# Rotavirus NSP1 activates human NLRP1 inflammasome

**DOI:** 10.64898/2026.10.26.761966

**Authors:** Qiru Zeng, Jinyi Sun, Stefano Bonazza, Debajyoti Chakraborty, Alexander Borodavka, Siyuan Ding

## Abstract

While inflammasomes detect and defend against pathogens, the precise mechanisms by which enteric viruses trigger these complexes, and the specific viral components involved remain incompletely understood. In this study, we demonstrate that rotavirus infection robustly induced human NLRP1 inflammasome activation via p38 mitogen-activated protein kinase (MAPK)-dependent and -independent pathways. Utilizing cell lines reconstituted with NLRP1 signaling and reverse genetics-rescued recombinant rotaviruses, we identified non-structural protein 1 (NSP1), a well-characterized viral antagonist of host interferon signaling, as an activator of the NLRP1 inflammasome in the absence of double-stranded RNA. Domain mapping analysis revealed that the C-terminus of NSP1 but not the C-terminal IRF3-binding site was essential for triggering inflammasome activation. Mechanistically, we show that NSP1 induced the proteasomal degradation of NLRP1 to elicit its activation, a process dependent on host neddylation pathways and the Cullin-3/Elongin B/C ubiquitin ligase complex. Together, these findings suggest that RV engages parallel NLRP1 surveillance pathways, including a p38-dependent response to infection, and a novel NSP1-dependent pathway involving the ubiquitin-proteosome system, where a viral strategy aimed at host protein degradation inadvertently trips the human NLRP1 inflammasome wire.

**SIGNIFICANCE STATEMENT:** When rotavirus infects the gut, it uses a viral protein called NSP1 to shut down key immune defenses of host cells. However, human cells have an immune sensor called NLRP1 that acts as a tripwire. When NSP1 attempts to force the destruction of NLRP1, the disposal process itself accidentally triggers the NLRP1 alarm. This causes an explosive inflammatory cell death that alerts the immune system. This discovery reveals how a viral strategy to suppress host immunity backfires, offering new insights into viral gut inflammation and future vaccine design.

## INTRODUCTION

Rotavirus (RV) is one of the leading causes of gastrointestinal diseases in young children worldwide. Non-structural-protein 1 (NSP1) plays a major role in antagonizing the host innate immune response by mediating the degradation of critical interferon (IFN) related genes such as IRF3, β-TrCP, and SAMD9 (1), (2), (3). While NSP1 is not required *in vitro*, it is critical for viral replication and pathogenesis in neonatal mice (4), presumably by antagonizing IFN signaling. Although less well-characterized, RV infection can also trigger Nlrp9b inflammasome activation and induce cleavage of gasdermin D, which leads to pyroptosis of infected intestinal epithelial cells (5). NLRP9 belongs to the NOD-like (NLR) family of cytosolic pattern recognition receptors, which are characterized by a central nucleotide-binding domain and C-terminal leucine-rich repeats. Several members of this family can initiate inflammasome assembly upon sensing pathogen- or damage-associated signals across mammalian and prokaryotic kingdoms (6).

NLRP1 is another member of this family and has emerged as a versatile innate immune sensor capable of responding to diverse stimuli. In addition to recognizing cytosolic double-stranded RNA (dsRNA) molecules (7), NLRP1 can be activated by pathogen-encoded proteases, toxins, and E3 ubiquitin ligases (8, 9) as well as ribotoxic stress responses via p38 MAPK signaling (10). These findings suggest that NLRP1 serves as an important hub for detecting pathogen-induced perturbations of cellular homeostasis. However, whether RV infection activates the NLRP1 inflammasome and, if so, which viral factor is responsible for this activity remain unknown. Here, we identify simian RV SA11 NSP1 as a potent activator of the human NLRP1 inflammasome and define a previously unrecognized role for this multifunctional viral immune modulator.

## RESULTS

### RV infection activates NLRP1 inflammasome

Although NLRP6 and NLRP9 inflammasomes are known to be activated by cytosolic dsRNA in the intestinal epithelium (5, 11), the functional roles of NLRP proteins have not been extensively characterized in the context of enteric viral infections. We reanalyzed our recent single-cell RNA sequencing dataset generated from mice in an experimental setting with or without RV infection (12). We found that while NLRP1B, NLRP3, and NLRP12 were highly expressed in immune cells of small intestines in mock-infected mice, NLRP1B/3/6/9B/10/12 were all expressed in intestinal epithelial cells post-infection (**Fig. 1A**). Given their expression patterns following RV infection, we selected these NLRPs for subsequent studies. We first generated HEK293 cells stably expressing ASC-GFP and then transfected them with NLRP constructs followed by RV infection to visualize speck formation by fluorescent microscopy as an indicator of inflammasome activation. With the exception of NLRP9, most NLRP proteins were expressed at comparable levels with and without RV infection (**Fig. 1B**). Transfection of NLRP1 and NLRP6 followed by RV infection led to robust ASC speck formation (**Fig. 1C and S1**). Notably, ASC-GFP, initially presented as cytosolic signals under mock conditions, formed bright punctate structures post NLRP inflammasome activation (**Fig. S1**). We further generated NLRP1 mutants where all five lysines in the Pyrin domain were replaced by arginines (PYD mut) or where a point mutation (D1401R) prevents oligomerization of the CARD domain (13) (**Fig. 1D**). While RV infectivity was similar in HEK293 cell lines expressing ASC-GFP and different NLRP1 constructs (**Fig. 1E**), RV infection induced ASC activation in up to 20% of cells transfected only with full-length NLRP1, whereas activation in cells expressing the NLRP1 mutants remained less than 8% (**Fig. 1F**). Val-boroPro (VbP), a DPP8/9 inhibitor (14), served as a positive control for NLRP1-mediated ASC activation (**Fig. 1F**). Taken together, our findings highlight that RV infection induces NLRP1 activation that requires intact PYD and CARD domain function.

**Figure 1.**
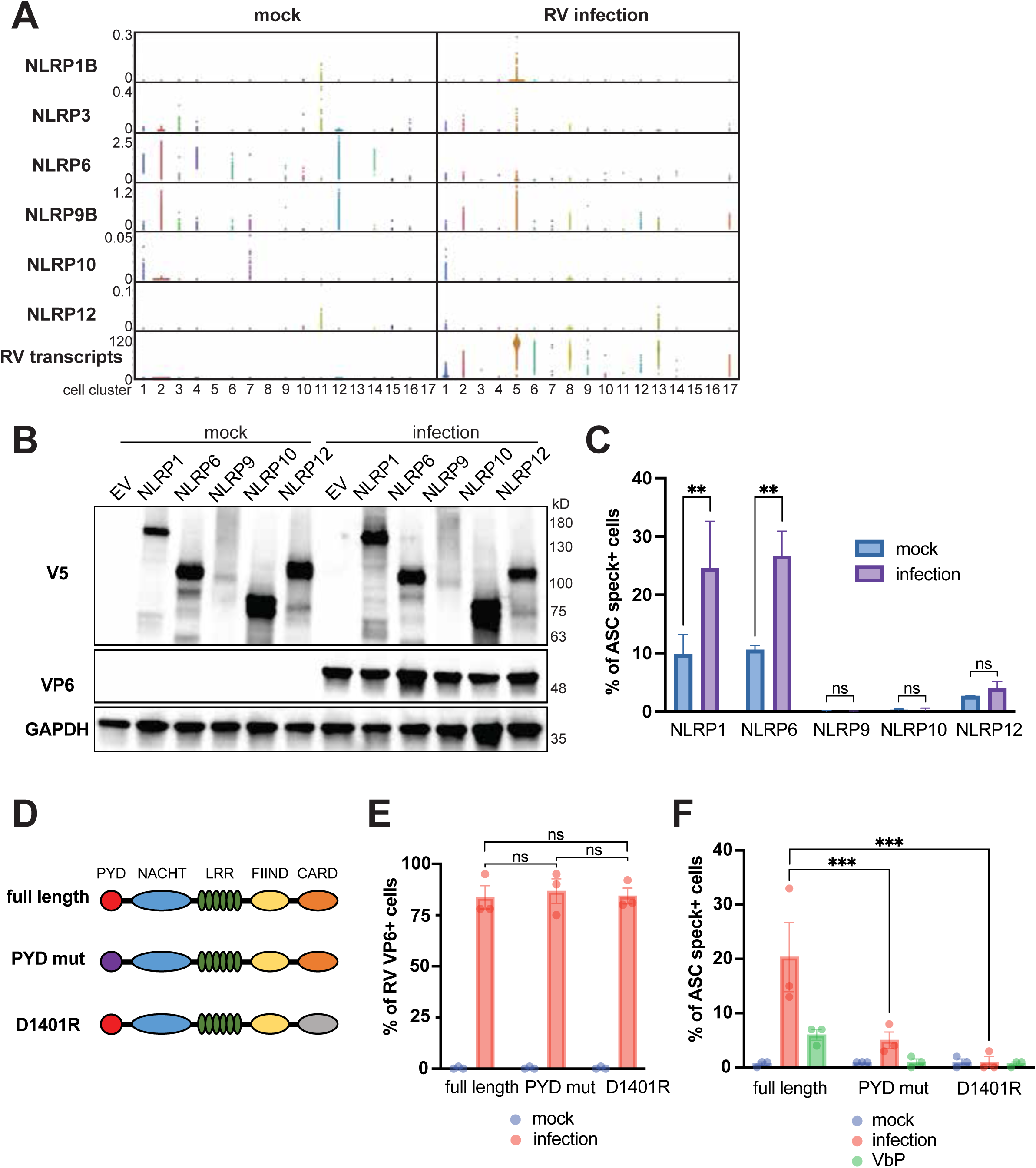
NLRP1 inflammasome is activated by rotavirus infection. (A) Transcript levels of NLRP genes for different clusters of intestinal epithelial cell subsets isolated from 5-day-old 129sv mice based on single-cell gene expression profiles. Immune cells: cluster 11 and 15; intestinal epithelial cells: cluster 1, 2, 3, 4, 5, 6, 7, 8, 9, 10, 12, 13, 14, 16, and 17. (B) Western blot confirming expression of V5-tagged NLRP constructs in HEK293-ASC-GFP cells with or without RV infection (SA11-4F strain, MOI=20, 8 hpi). (C) Quantification of ASC activation in HEK293-ASC-GFP cells transfected with NLRP1, 6, 9, 10 or 12 with or without SA11-4F infection (MOI=20, 8 hpi), shown as percentage of speck-positive cells. Each bar represents 2 biological repeats. (D) Schematic of NLRP1 mutant design. The indicated mutations were introduced to NLRP1 sequences for subsequent generation of mutant constructs. (E-F) Flow cytometry analysis of ASC activation in HEK293-ASC-GFP cells transfected with indicated NLRP1 mutants followed by RV infection (MOI=1, 24hpi). (E) RV infection levels, quantified as the percentage of VP6-positive cells. (F) ASC speck formation, quantified as the percentage of ASC-speck-positive cells.

### SA11 NSP1 mediates NLRP1 activation by ectopic expression or during RV infection

To extend our investigation to the viral determinants of NLRP1 activation, we infected HEK293 cells expressing ASC-GFP and NLRP1 with RV at high multiplicity of infection (MOI) or low MOI. We observed ASC activation in up to ∼75% or ∼55% of cells, respectively, quantified by flow cytometry (**Fig. S2A and B**). Treatment with SB202190, a p38 MAPK inhibitor (10), had no effect on this RV-induced ASC activation (**Fig. S2A and B**), suggesting that the RV-induced NLRP1 inflammasome activation in these cells was likely independent of p38 MAPK signaling. Therefore, we hypothesized that newly synthesized viral proteins, instead of the incoming dsRNA genome, serve as the trigger for NLRP1 inflammasome activation. Specifically, we examined NSP1 due to its established role in targeting cellular substrates for degradation. Using known NLRP1 activators such as human rhinovirus 3C protease (15) and TAOK2 catalytic domain (10) as positive controls, we found that ectopic expression of Flag-tagged NSP1 derived from the simian RV SA11 strain was able to induce comparable levels of activation (**Fig. 2A and B**). NSP1 is among the most sequence-divergent RV proteins, sharing as little as 36% amino-acid identity between some mammalian RV strains, with simian and murine NSP1 proteins clustering more closely together (**Fig. 2C**). Intriguingly, while NSP1 expression levels were similar among various RV strains, as measured by the percentage of Flag-positive cells by flow cytometry (**Fig. 2D**), only SA11 NSP1 successfully induced ASC activation in up to ∼60% of cells (**Fig. 2E**). To further validate this finding in the context of viral infection, we employed optimized plasmid-based RV reverse genetics (16) to create recombinant RVs, including parental simian SA11 strain with its native NSP1 replaced by that derived from simian RRV strain, human DS-1 strain, and bovine RV strain (**Fig. 2F**). We also generated the reciprocal monoreassortant RVs by replacing the native bovine RV RF strain NSP1 with SA11 NSP1 in the RF backbone (**Fig. 2F**). Remarkably, in both conditions, SA11 NSP1 conferred strong NLRP1 inflammasome activation (**Fig. 2G and H**), regardless of the RV genetic background. In addition to HEK293 cells, we also tested A549 cells stably reconstituted with both ASC-GFP and NLRP1. RV infection induced NLRP1 activation in an MOI-dependent manner, with ASC specks visualized in viral antigen VP6 positive cells (**Fig. S2C and D**). This activation was dependent on both NSP1 and p38 MAPK pathway (**Fig. S2E and F**). A recombinant NSP1-deletion virus that is fully replication competent *in vitro* (17) induced NLRP1 activation, which was completely blocked by SB202190 (**Fig. S2E and F**), indicating that RV can activate NLRP1 through both an NSP1-dependent pathway, and an NSP1-independent, p38-dependent pathway, with their relative contributions varying between cell types. These observations confirm that SA11 NSP1 expression individually, or during the course of RV infection, is sufficient to induce NLRP1 activation.

**Figure 2.**
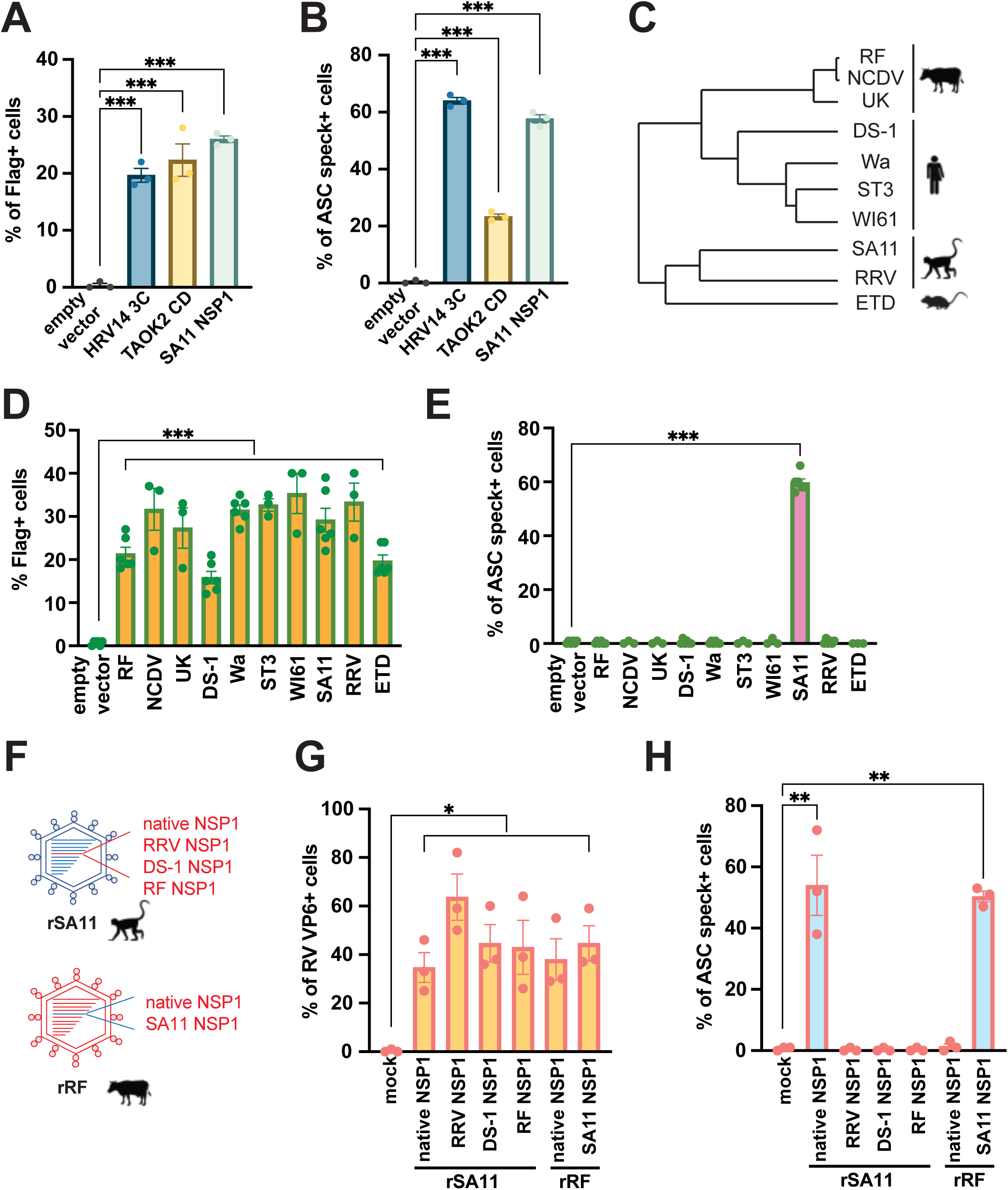
NLRP1 inflammasome is activated by RV NSP1. (A-B) Flow cytometry analysis of ASC activation in HEK293-ASC-GFP cells co-transfected with 0.5 μg of wild-type NLRP1 and 0.5 μg of Flag-tagged empty vector, human rhinovirus 3C protease (HRV14 3C), TAOK2 catalytic domain (TAOK2 CD), or SA11 NSP1, for 24hrs. (A) Ectopic expression of indicated Flag-tagged constructs of NLRP1 activators, quantified as the percentage of Flag-positive cells. (B) ASC speck formation, quantified as the percentage of ASC-speck-positive cells. (C) Phylogenetic tree of rotavirus strains employed in this study. (D-E) Flow cytometry analysis of ASC activation in HEK293-ASC-GFP cells co-transfected with 0.5 μg of wild-type NLRP1 and 0.5 μg of Flag-tagged empty vector or indicated RV NSP1s, for 24hrs. (D) Ectopic expression of indicated Flag-tagged constructs, quantified as the percentage of Flag-positive cells. (E) ASC speck formation, quantified as the percentage of ASC-speck-positive cells. (F) Schematic of reverse genetics strategy used to generate recombinant RV strains with NSP1 from different strains in the SA11 backbone, as well as SA11 or native NSP1 in the RF backbone. (G-H) Flow cytometry analysis of ASC activation in HEK293-ASC-GFP cells transfected with wild-type NLRP1 followed by infection with indicated recombinant RVs (MOI=1, 24hpi). (G) RV infection levels, quantified as the percentage of VP6-positive cells. (H) ASC speck formation, quantified as the percentage of ASC-speck-positive cells.

### C-terminus of SA11 NSP1 is required for association with NLRP1

We next sought to investigate the molecular mechanisms underlying SA11 NSP1-mediated NLRP1 activation. To this end, we utilized naturally occurring SA11 variants including SA11-5S, which contains a C-terminal 17-amino-acid truncation important for IRF3 binding (18), and SA11-30-1A, which contains a C-terminal 71-amino-acid truncation (19) (**Fig. 3A**). A549-ASC-NLRP1 cells were infected with the parental SA11-4F, SA11-5S variant, or SA11-30-1A variant, and ASC-GFP speck formation was assessed at 8 hours post-infection (**Fig. 3B and S3**). Interestingly, the extent of ASC activation correlated with the length of the C-terminal truncation. Whereas parental SA11-4F strain induced ASC activation in ∼50% of cells, this response triggered by SA11-5S was reduced to only ∼30% of cells, and was barely detectable with SA11-30-1A (**Fig. 3B**), indicating that the C-terminus contributes to infectivity and SA11 NSP1-induced ASC activation. In addition to natural viral variants, we introduced a series of mutations in the N-terminal RING-finger domain of SA11 NSP1, including key cysteine and histidine residues H61R/C63A/C66A and C42A (2), and the C-terminal end (**Fig. 3C**), and studied their effects through ectopic expression. With comparable NSP1 expression levels (**Fig. 3D**), WT full-length NSP1 induced ASC activation in nearly 60% of cells (**Fig. 3E**). In contrast, the proportion of cells exhibiting ASC specks was reduced to ∼40% following expression of SA11-NSP1 containing a C-terminal 17-amino-acid truncation (**Fig. 3E**), consistent with the phenotype observed in SA11-5S-infected cells (**Fig. 3B**). Furthermore, minimal ASC speck formation was observed in cells following expression of SA11-NSP1 containing a C-terminal 42-amino-acid-truncation, the RING domain mutant, and a C42A mutant with the zinc-binding site disrupted (**Fig. 3E**). Using GFP-tagged NSP1 and GFP-antibody conjugated beads, we could co-precipitate V5-tagged NLRP1 in transfected cells (**Fig. 3F**). We also performed the reciprocal immunoprecipitation experiments in which V5-tagged NLRP1 and V5-antibodies co-precipitated full-length WT SA11-4F NSP1 but not deletion mutants that are truncated by 17, 42, and 71 amino acids (**Fig. 3G**). Together, these data support that the C-terminal region of NSP1 mediates its interaction with NLRP1 and is important for efficient NLRP1 inflammasome activation.

**Figure 3.**
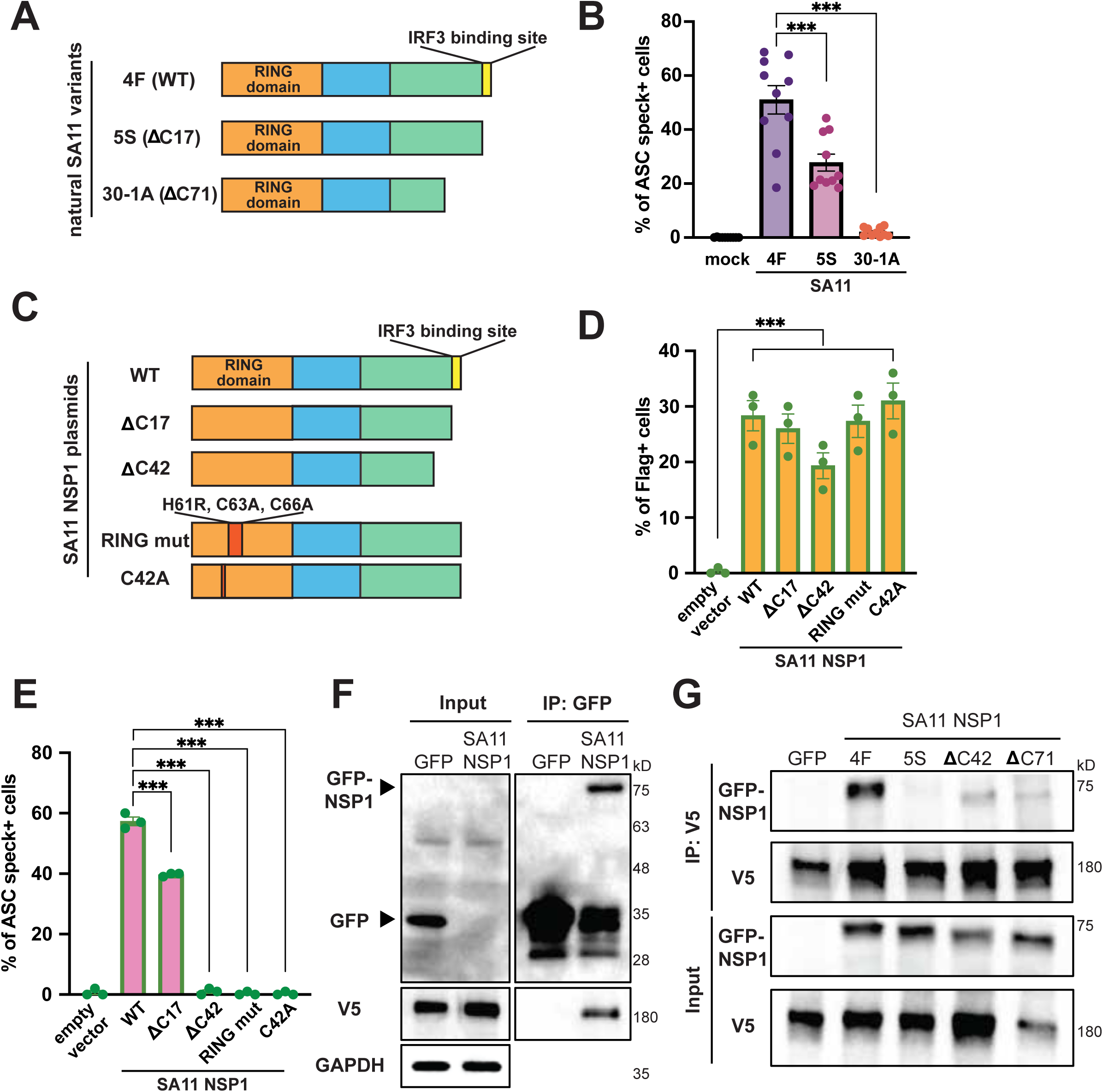
NLRP1 interacts with SA11-4F-NSP1 on its C-terminus. (A) Schematics of SA11 natural variants used in this study. (B) Quantification of ASC activation in A549-ASC-GFP-NLRP1 cells infected with indicated SA11 natural variants (MOI=10, 8 hpi), shown as percentage of ASC-speck-positive cells. (C) Schematics of SA11 mutant constructs used in this study, including progressive C-terminal truncations and mutations in the RING domain. (D-E) Flow cytometry analysis of ASC activation in HEK293-ASC-GFP cells co-transfected with 0.5 μg of wild-type NLRP1 and 0.5 μg of Flag-tagged SA11 NSP1 or its corresponding mutants for 24hr. (D) Ectopic expression of indicated Flag-tagged constructs, quantified as the percentage of Flag-positive cells. (E) ASC speck formation, quantified as the percentage of ASC-speck-positive cells. (F) Lysates of HEK293 cells transfected with tagRFP-NLRP1-V5 and pG-LAP6 empty vector or SA11-4F-NSP1 for 48 hours were subject to immunoprecipitation using GFP-trap beads and probed for GFP, V5 and GAPDH. (G) Lysates of HEK293-ASC cells co-transfected with tagRFP-NLRP1-V5 and pG-LAP6 empty vector, or SA11-4F-NSP1, or NSP1 mutants for 48 hours were subject to immunoprecipitation using V5-conjugated beads and probed for GFP, V5, and GAPDH.

### NSP1 as a viral E3 ligase mediates NLRP1 degradation and activation

Because NSP1 targets cellular substrates for degradation (20) and bacterial E3 ligases can activate NLRP1, we hypothesized that SA11 NSP1 may act as a viral E3 ligase to promote NLRP1 degradation and activation. We directly tested this hypothesis by examining NLRP1 degradation in RV infected A549-ASC-NLRP1 cells and treating infected cells with MLN4924, a neddylation inhibitor (21). Given the dual activation pathways identified in A549 cells, we included SB202190 treatment to exclude effects mediated through the MAPK-dependent pathway. Indeed, NLRP1 protein levels were substantially reduced by RV infection and were partially restored by MLN4924 but not SB202190 treatment, suggesting that RV infection-induced NLRP1 degradation in A549 cells is independent of the p38 MAPK signaling (**Fig. 4A**). Consistent with the above findings, we observed that treatment with MG132, a broad-spectrum 26S proteasome inhibitor, and TAK-243, a ubiquitination E1 ligase inhibitor (22), prevented RV infection-induced NLRP1 activation (**Fig. 4B and S4A**). A similar phenotype was observed in HEK293 cells where ASC activation by RV infection under inhibitor treatment was analyzed by flow cytometry (**Fig. 4C and 4D**). Treatment of MG132, Bortezomib, another proteasome inhibitor, MLN4924, or TAK-243 efficiently suppressed ASC activation by RV infection (**Fig. 4D**). These data indicate that RV-induced ASC activation relies on proteasome and neddylation pathways. NSP1 has recently been shown to hijack the host Cullin-3 (CUL3) and Elongin B/C E3 ubiquitin ligase complex to target cellular substrates for degradation (2, 23). To determine whether these activities contribute to NLRP1 activation, we utilized siRNA to transiently knock down *TCEB1 and TCEB2* which encode elongin C and elongin B, respectively, or *CUL3* in A549-ASC-NLRP1 cells and infected with RV. We observed NLRP1 degradation was largely abolished when CUL3 was knocked down (**Fig. 4E**). In parallel, we utilized recombinant RVs carrying NSP1 mutants that disrupt interactions with ELOC or CUL3 (20) to test the functional impact on NLRP1 activation. Compared to the parental SA11 and the IRF3-binding mutant (NSP1^*IRF3^), which produced comparable levels of ASC speck formation, ELOC mutant (NSP1^*ELOC^) or CUL3 mutant (NSP1^*CUL3^) induced modestly reduced ASC specks in infected A549 cells in the presence of SB202190 (**Fig. 4F and S4B**). Taken together, these data suggest that RV NSP1 inadvertently activates NLRP1 inflammasome due to its ability to hijack the host CUL3 E3 ligase complex (**Fig. 4G**).

**Figure 4.**
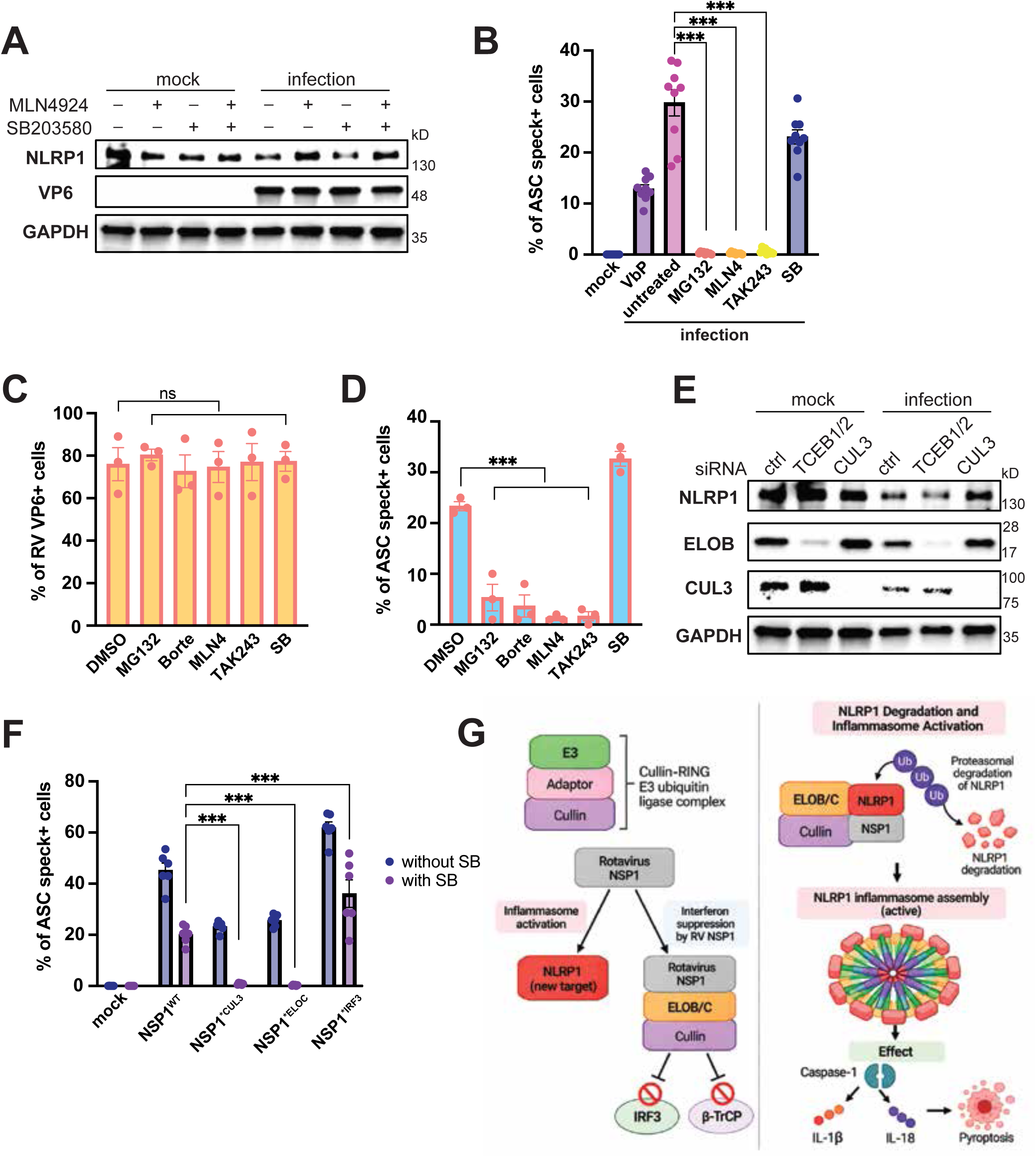
SA11-4F-NSP1 regulates NLRP1 degradation and activation. (A) Western blot results of A549-ASC-GFP-NLRP1 cells infected with SA11-4F an MOI of 10 with or without treatment of SB202190 (20μM) or MLN4924 (10μM) post-inoculation. Cell lysates were harvest at 24hpi and probed for NLRP1, VP6, and GAPDH. (B) Quantification of ASC activation in A549-ASC-GFP-NLRP1 cells infected with SA11 (MOI=10, 8hpi), with or without treatment of SB202190 (20μM), MG132 (25μM), MLN4924 (10μM) or TAK-243 (5μM) post-inoculation, shown as percentage of speck-positive cells. Each bar represents 3 biological repeats. VbP (30μM) treatment served as a control for ASC activation. (C-D) Flow cytometry analysis of ASC activation in HEK293-ASC-GFP cells transfected with wild-type NLRP1 followed by infection with RV (MOI=1, 24hpi), treated with DMSO, MLN4924 (10μM), TAK-243 (5μM), bortezomib (10μM), MG132 (10μM), or SB202190 (20μM) post-innoculation. (C) RV infection levels, quantified as the percentage of VP6-positive cells. (D) ASC speck formation, quantified as the percentage of ASC-speck-positive cells. (E) Western blot results of A549-ASC-GFP-NLRP1 cells transfected with control, *TCEB1/2*, or *CUL3* siRNA, left uninfected (mock) or infected with SA11-4F (MOI=10, 24hpi). Cell lysates were probed for NLRP1, TCEB2, CUL3, and GAPDH. (F) Quantification of ASC activation in A549-ASC-GFP-NLRP1 cells infected with SA11-4F and its corresponding mutants SA11-4F-NSP1*^CUL3^, SA11-4F-NSP1*^ELOC^, or SA11-4F-NSP1*^IRF3^ (MOI=10, 8hpi), with or without treatment of SB202190 (20μM) 1h pre-inoculation, shown as percentage of speck-positive cells. Each bar represents 2 biological repeats. (G) A working model depicting the mechanism of NLRP1 degradation and activation by NSP1 based on the findings in this study.

## Discussion

In this study, we identify rotavirus NSP1 as a previously unrecognized activator of the human NLRP1 inflammasome. Using both ectopic expression and recombinant RVs generated by reverse genetics, we demonstrate that NSP1 is sufficient for NLRP1 activation. Whereas A549 cells exhibited both NSP1-dependent and MAPK-dependent pathways, NLRP1 activation in HEK293 cells was independent of p38 MAPK signaling, a pathway previously implicated in NLRP1 activation during ribotoxic stress responses. Interestingly, the residual NLRP1 activation observed following infection with the NSP1-deletion virus was sensitive to p38 inhibition, suggesting that rotavirus can engage at least two distinct NLRP1 activation pathways. Thus, NLRP1 may provide parallel surveillance of rotavirus infection by sensing infection-associated stress through p38 signaling while independently responding to NSP1-mediated perturbation of the ubiquitin-proteasome system. The relative contribution of these pathways appears to be cell-context dependent. Instead, NLRP1 activation in HEK293 cells required NSP1 and relied on ubiquitin-proteasome system and neddylation machinery. Together, these findings support a model in which NSP1 functions as a viral E3 ligase that targets NLRP1 for degradation, thereby triggering inflammasome activation.

Our findings further highlight the remarkable functional diversity of NSP1 proteins among RV strains. Although NSP1 proteins are highly divergent and share a common role as innate immune antagonists, only SA11 NSP1 robustly activated NLRP1. This observation parallels previous studies showing that SA11 NSP1 possesses unique substrate specificities, *i.e.*, RNase L degradation, that are not shared by even highly similar simian RV RRV strain (24). The ability of SA11 NSP1 to activate NLRP1 therefore adds another distinctive activity to its functional repertoire and suggests that strain-specific adaptations have expanded the range of host proteins recognized by this viral factor.

NSP1 has evolved multiple mechanisms to engage distinct cellular substrates. Previous studies have shown that different regions of NSP1 mediate interactions with IRF proteins, β-TrCP, RNase L, SAMD9, and additional innate immune factors. In many cases, substrate recognition is determined by variable motifs located within the C-terminal region of NSP1, whereas the conserved N-terminal RING domain recruits components of the host ubiquitination machinery to promote substrate degradation. Consistent with this paradigm, our mutagenesis studies identified both the RING domain and a discrete C-terminal region of SA11 NSP1 as critical for NLRP1 activation. These findings suggest that NLRP1 may represent a *bona fide* NSP1 substrate whose recognition requires specialized determinants present uniquely in SA11 NSP1. The strain-specific ability of SA11 NSP1 to activate NLRP1 may therefore reflect the distinct evolutionary histories of NSP1 among these viruses, as whole-genome analyses concluded that RRV, TUCH, PTRV, and SA11 arose through different reassortment events that do not share common evolutionary origins for many genome segments, including NSP1.

The observation that proteasome, ubiquitination, and neddylation inhibitors all suppressed NLRP1 activation provides additional support for a degradation-based mechanism. Multiple NLRP1 activation pathways converge on proteasome-mediated destruction of inhibitory N-terminal regions, including activation by pathogen-derived enzymes and cellular E3 ligases. Our data suggest that NSP1 exploits a similar mechanism, either through direct ubiquitination of NLRP1 or through modification of a closely associated regulatory factor. Future studies will be needed to define the precise ubiquitination sites, determine whether NLRP1 is a direct substrate of NSP1, and establish the molecular details of the binding interface. The availability of Nlrp1b-deficient mice, together with recombinant murine RVs engineered to express SA11 NSP1, provides a unique opportunity to investigate the physiological significance of this pathway *in vivo*. Such studies should enable direct assessment of how NLRP1 activation influences viral replication, epithelial cell death, intestinal inflammation, and disease severity during RV infection.

More broadly, our findings reveal an intriguing duality in NSP1 function. NSP1 is best known as a potent antagonist of innate immunity, targeting multiple antiviral pathways to promote viral replication. However, our results demonstrate that the same viral protein can also activate an inflammatory cell death pathway. Whether NLRP1 activation ultimately benefits the host by restricting RV infection or benefits the virus through tissue damage, epithelial turnover, or modulation of local immune responses remains unclear. Our findings raise the possibility that excessive inflammasome activation in the intestinal epithelium may contribute to pathological outcomes associated with RV infection or vaccination. In particular, it will be interesting to determine whether NLRP1 activation contributes to intestinal inflammation, altered epithelial barrier function, or dysregulated tissue remodeling processes linked to intussusception, a rare but serious adverse event associated with certain live attenuated RV vaccines. Although there is currently no direct evidence connecting NLRP1 signaling to vaccine-associated intussusception, the identification of NSP1 as a potent NLRP1 activator provides a mechanistic framework to explore whether strain-specific differences in inflammasome activation influence intestinal pathology in susceptible hosts.

## ACKNOWLEDGEMENT

This study is supported by the National Institutes of Health (NIH) grant R01 AI150796 and Gates Foundation grant INV-075616 to S.D. The funders had no role in study design, data collection and analysis, decision to publish, or preparation of the manuscript. We thank all members of the Ding lab for helpful discussion. A.B. acknowledges funding from Wellcome Trust (307249/Z/23/Z). We also thank Drs. Patrick Mitchell (University of Washington) and Andrew Sandstrom (University of Texas Southwestern Medical Center) for discussing unpublished data.

## MATERIALS AND METHODS

### Reagents, cells, and viruses

#### Reagents

SB202190 (Selleckchem, S1077), VbP (Selleckchem, S8455), MG132 (Selleckchem, S2619), MLN4924 (Selleckchem, S7109), TAK-243 (Selleckchem, S8341), Bortezomib (Selleckchem, S1013), and Normocin (Invivogen, ant-nr-1) were purchased commercially.

#### Cells

Wild-type HEK293 cells were cultured in DMEM (Gibco, 12491050) supplemented with 10% heat-inactivated fetal bovine serum (VWR, 89510-186) and 1X Penicillin-Streptomycin-Glutamine (Thermo Scientific, 10378016). Reconstituted HEK293-ASC-GFP cells were generated by transfection followed by single-cell sorting and clonal expansion. Wild-type HEK293 cells were transfected with pLEX-MCS-ASC-GFP (Addgene #73957), and GFP-positive singlets were sorted into individual wells in 96-well plates by BD FACSAria II Cell Sorters at the Flow Cytometry & Fluorescence Activated Cell Sorting Core at Washington University. Single-cell clones were grown until visible colonies formed, and then expanded for experimental use. Wild-type and MA104 cells (ATCC, CRL-2378.1) were grown in M199 (Thermo Scientific, 11150067) supplemented with 10% heat-inactivated fetal bovine serum (VWR, 89510-186) and 1X Penicillin-Streptomycin-Glutamine (Thermo Scientific, 10378016). A549-ASC-GFP-NLRP1 cells (Invivogen, a549-ascg-nlrp1-av) were cultured in DMEM supplemented with 10% heat-inactivated fetal bovine serum (VWR, 89510-186), 1X Penicillin-Streptomycin-Glutamine (Thermo Scientific, 10378016), and 100 μg/mL of normocin. The ASC-GFP expression in the A549-ASC-GFP-NLRP1 cells was manually inducted by treatment of recombinant human TNF-α (R&D Systems, 410-MT-025/CF) at 5 ng/mL overnight.

#### Viruses

Simian RVs, including SA11 natural variants and mutants, SA11 NSP1 deletion virus (17), and bovine RVs were cultured and titrated in MA104 cells in serum-free M199 medium. Prior to infection, RVs were activated by incubating with 5μg/mL of trypsin at 37°C for 25 minutes. Cells were washed with serum-free medium twice and were inoculated with activated RVs at desired MOIs at 37°C for 1 hour. Following absorption, the inoculum was then removed and replaced with complete medium to minimize cell damage from prolonged serum deprivation.

#### Plasmids

NLRP1 in pLenti6.3/V5-DEST (DNASU, HsCD00860948), NLRP6 in pLenti6.3/V5-DEST (DNASU, HsCD00954494), NLRP9 in pLenti6.3/V5-DEST (DNASU, HsCD00852567), NLRP10 in pLenti6.3/V5-DEST (DNASU, HsCD00852857), and NLRP12 in pLenti6.3/V5-DEST (DNASU, HsCD00852096) were purchased commercially. NLRP1 PYD mutant (K12R/K15R/K41R/K54R/K58R) and NLRP1 D1401R mutant are used as previously constructed (10). pLEX-MCS-ASC-GFP (Addgene #73957) was used for HEK293-ASC-GFP generation. V5-tagged NLRP1 construct was generated by Gateway cloning using pDONR221-NLRP1 (DNASU HsCD00860481) as the entry clone and pDEST pEF1alpha FRT Tag-eRFP as the destination vector. pG-LAP6 empty vector (EV) and pG-LAP6-SA11-5S-NSP1 were previously reported (2), pG-LAP6-SA11-4F-NSP1 and mutants (pG-LAP6-SA11-4F-NSP1-C42-deletion, pG-LAP6-SA11-4F-NSP1-C71-deletion, pG-LAP6-SA11-4F-NSP1-RING-mutant, and pG-LAP6-SA11-4F-NSP1-C42A) were generated by Genscript.

#### Single-cell RNA sequencing

Single-cell RNA sequencing was performed as described previously (25). Briefly, 5-day-old 129Sv mice were orally inoculated with 5×10^6^ FFU of murine rotavirus D6/2 strain. At 12 hpi, small intestinal cells were isolated using an EDTA-DTT extraction protocol and sorted as previously described. Live cells were purified using a BD FACS Aria II flow cytometer. Single-cell library preparation and sequencing were performed at the Stanford Genome Sequencing Service Center (GSSC). Libraries were prepared using the Chromium Single Cell 3′ Library and Gel Bead Kit v2 (10× Genomics, PN-120237) according to the manufacturer’s instructions. Libraries were sequenced on an Illumina NextSeq 500 platform using a 150-cycle High Output V2 kit with the following configuration: Read 1, 26 cycles; Read 2, 98 cycles; Index 1, 8 cycles. Single-cell expression analysis was performed using the Seurat package. Viral gene expression scores were calculated using the AddModuleScore function.

#### Plasmid and siRNA transfection

Plasmids were transfected at 1μg per well in 24-well plates and 5μg per well in 6-well plates in HEK293 cells using Lipofectamine™ 3000 (Invitrogen, L3000015) per manufacturer’s instructions. Scrambled, *TCEB1*, *TCEB2,* and *Cullin-3* siRNAs (SMARTpool, Dharmacon) were transfected at 5 nM per well in 24-well plates using Lipofectamine® RNAiMAX (Invitrogen, 13778-075) per manufacturer’s instructions.

#### Western blot

Cells were washed with PBS and lysed by RIPA buffer (Thermo Scientific, 89901) supplemented with 100X protease inhibitor cocktail and phosphatase inhibitor (Thermo Scientific, 78420), followed by a 10-minute incubation on ice (26). Cell lysates were then subjected to centrifugation at 14,000 RPM for 10 minutes at 4°C to remove cell debris and chromatids. The protein samples were then boiled in 2X Laemmli Sample Buffer (Bio-Rad, #1610737EDU) containing 5% β-mercaptoethanol at 95 °C for 5 minutes. Prepared samples were run in 4-12% gels and transferred onto nitrocellulose membranes. Membranes were blocked in 5% BSA in TBS + 0.1% Tween-20 (TBST) at room temperature before incubation at 4°C overnight with primary antibodies: anti-V5 (Cell Signaling Technology, 80076), anti-GAPDH (Cell Signaling Technology, 2118L), anti-GFP (Santa Cruz, sc-9996), and anti-VP6 (Santa Cruz, sc101363).

#### Reverse Genetics

The rotavirus reverse genetics system was performed as previously described (27). Briefly, BHK-T7 cells were transfected with plasmids encoding the 11 rotavirus genome segments of either simian SA11 strain or bovine RF strain under the control of T7 promoters together with required helper plasmids. Sixteen hours after transfection, cells were washed and cultured in serum-free medium. Twenty-four hours later, *SERPINB1*-knockout MA104 cells were added to amplify rescued viruses. After 3 days, supernatants containing rescued viruses were collected and used to infect fresh MA104 cells. Virus rescue was confirmed by monitoring cytopathic effects and analyzing viral gene expression.

#### Immunoprecipitation

Cells were washed with PBS and lysed using NP40 buffer, followed by a 10-minute incubation on ice. Cell lysates were then subjected to centrifugation at 14,000 RPM for 10 minutes at 4°C to remove cell debris and chromatids. Then, 1/10 volume of each harvested protein sample was used as input, the rest was incubated overnight at 4°C with Dynabeads protein G conjugated with V5 (E9H8O) Mouse Monoclonal Antibody (Cell Signaling, 80076) following the manufacturer’s instructions. Next, using a magnet stand, the beads were washed three times with NP40 buffer. Then, the beads were resuspended in 2X Laemmli Sample Buffer, and boiled at 98°C for 10 minutes before western blot analysis.

#### Flow cytometry

To quantify the numbers of ASC-EGFP specks, cells were seeded in 24-well plates, treated with compound or infected with RV, and analyzed by flow cytometry ATTUNE NXT 4 LASER. Cells were not fixed and live cells were gated based on forward and side scatter for singlets, and with Zombie Aqua Live (Biolegend, 423102) for live cells. The results were analyzed by FlowJo 10.8.1 software.

#### Microscopy and quantification

The quantification of ASC speck formation was performed with a custom automated analysis pipeline. Multi-channel fluorescence images were acquired according to the relevant experiment and segmentation was carried out in a jupyter notebook with cellpose (28). The package employs pre-trained machine learning models to determine the cell boundaries from the DAPI channel and export them as FIJI ROIs. The model cpsam_v2 was used in this study. Subsequently, a FIJI script loads the masks and the FITC channel for all images, and performs the analysis, counting the ASC specks in each cell automatically for the whole dataset. Specifically, first masks at the edges of the FOV are filtered to remove cells only partially in view, and a background rolling ball filter is applied. Then, the script cycles through each cell and detects the presence of a speck by automatic thresholding. Each speck is then compiled and the number of positive and negative cells is calculated for each image, across conditions and replicates. The code used for this project is available on request: Segmentation.ipynb Analysis_extended.ijm

#### Statistical analysis

For Figures 1C, 1E, 1F, 4E, and S2F, two-way ANOVA and Šidák test was used; For Figures 2A, 2B, 2D, 2E, 2G, 2H, 3B, 3D, 3E, 4B, 4C, and 4D, one-way ANOVA and Dunnet test was used.

---

**Supplemental Figure 1.**
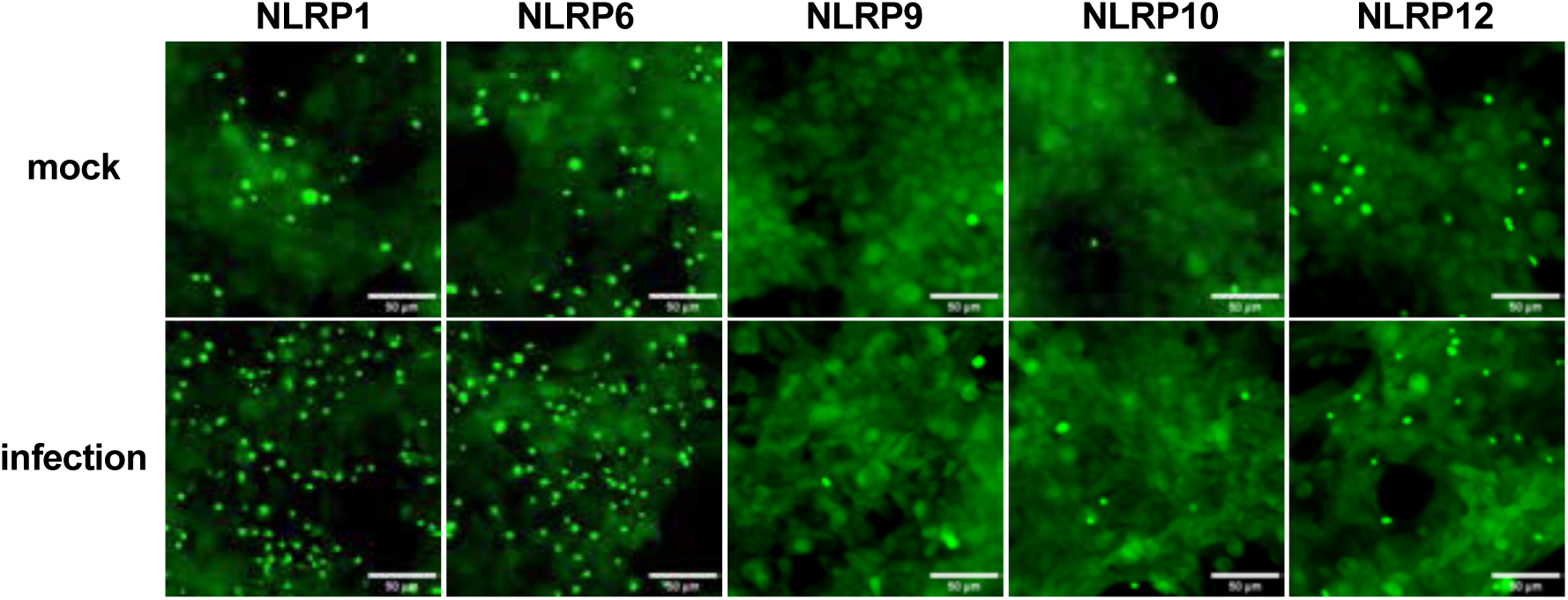
Representative fluorescence microscopy images of HEK293-ASC-GFP cells transfected with NLRP1, 6, 9, 10 or 12. Images show representative fields following mock treatment or RV infection (MOI=20, 8hpi). Scale bars, 50μm.

**Supplemental Figure 2.**
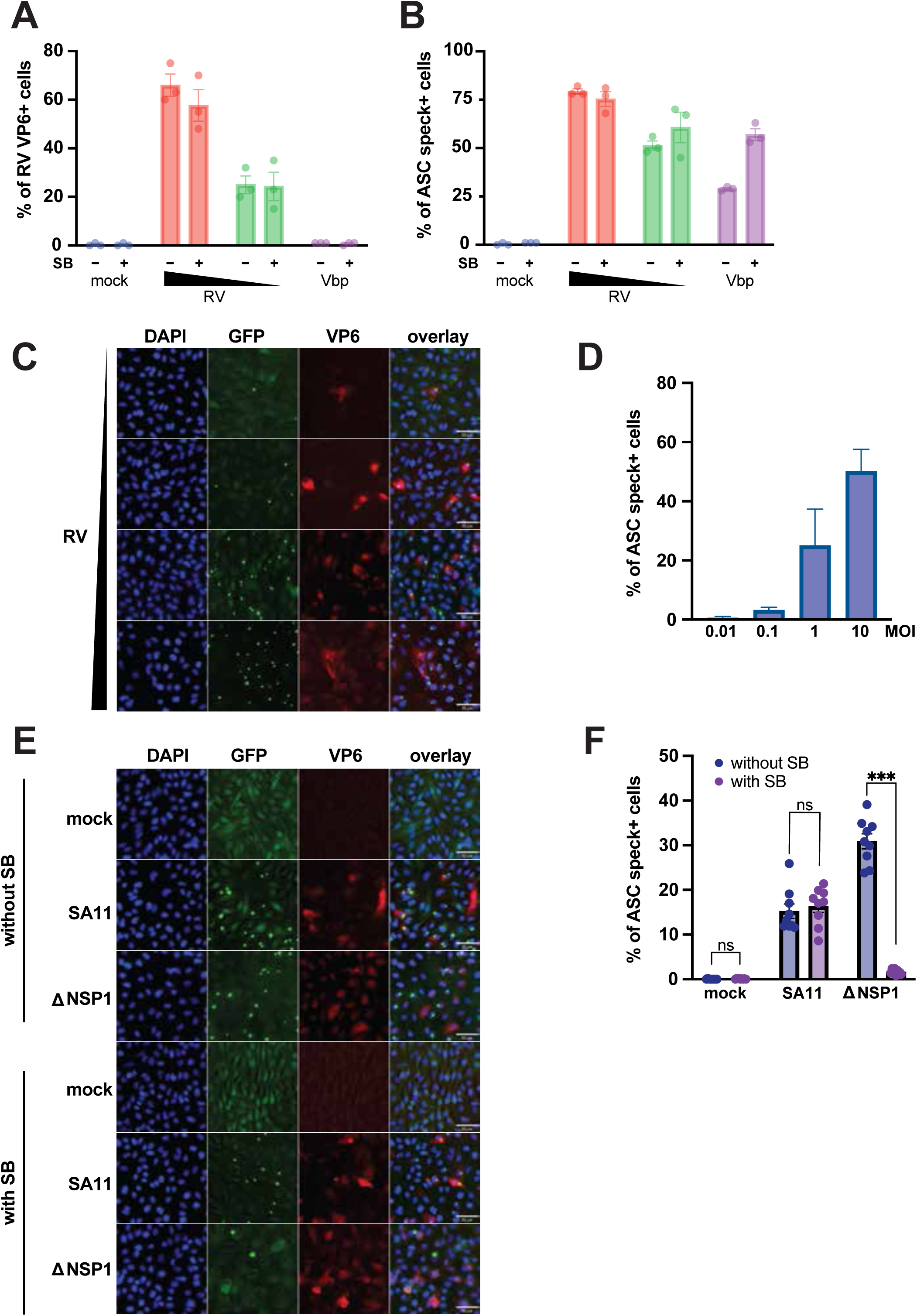
(A-B) Flow cytometry analysis of ASC activation in HEK293-ASC-GFP cells co-transfected with indicated NLRP1 constructs followed by RV infection (MOI=1, 24 hpi) with or without SB202190 (20μM), or VbP (30μM) treatment for 24h. (A) RV infection levels, quantified as the percentage of VP6-positive cells. (B) ASC speck formation, quantified as the percentage of ASC-speck-positive cells. (C) Representative fluorescence microscopy images of A549-ASC-GFP-NLRP1 cells infected with SA11-4F at MOIs of 0.01, 0.1, 1 or 10, at 8hpi. Images show representative fields. Scale bars, 50μm. (D) Quantification of ASC activation in A549-ASC-GFP-NLRP1 cells infected with SA11-4F at MOIs of 0.01, 0.1, 1 or 10, at 8 hpi, shown as percentage of speck-positive cells. Each bar represents 3 biological repeats. (E) Representative fluorescence microscopy images of A549-ASC-GFP-NLRP1 cells infected with SA11-4F or recombinant SA11-NSP1-deletion (MOI=10, 8hpi), shown as percentage of speck-positive cells. Each bar represents 3 biological repeats. (F) Quantification of ASC activation in A549-ASC-GFP-NLRP1 cells either mock or infected with SA11-4F or recombinant SA11-NSP1-deletion (MOI=10, 8hpi), shown as percentage of speck-positive cells. Each bar represents 3 biological repeats.

**Supplemental Figure 3.**
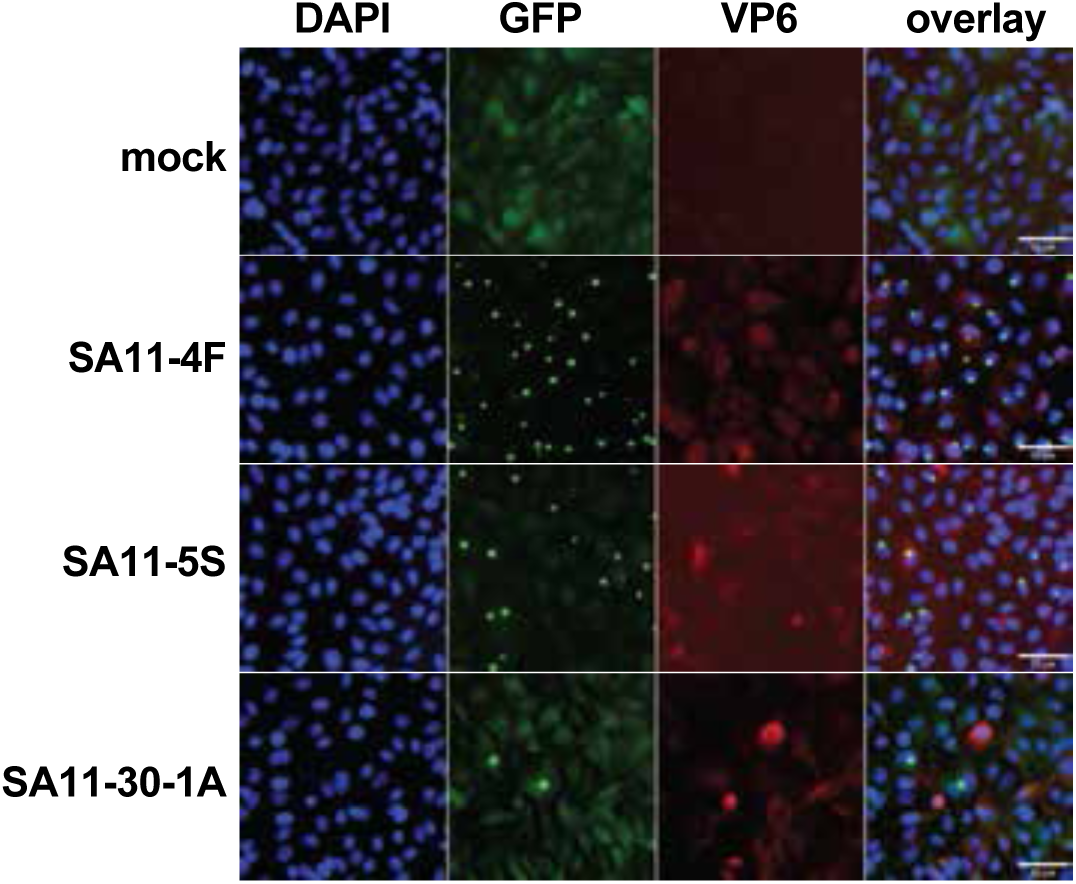
Representative fluorescence microscopy images of A549-ASC-GFP-NLRP1 cells infected with SA11 (MOI=10, 8hpi), with or without treatment of SB202190 (20μM), MG132 (25μM), MLN4924 (10μM) or TAK-243 (5μM) post-inoculation, shown as percentage of speck-positive cells. Images show representative fields. Scale bars, 50μm.

**Supplemental Figure 4.**
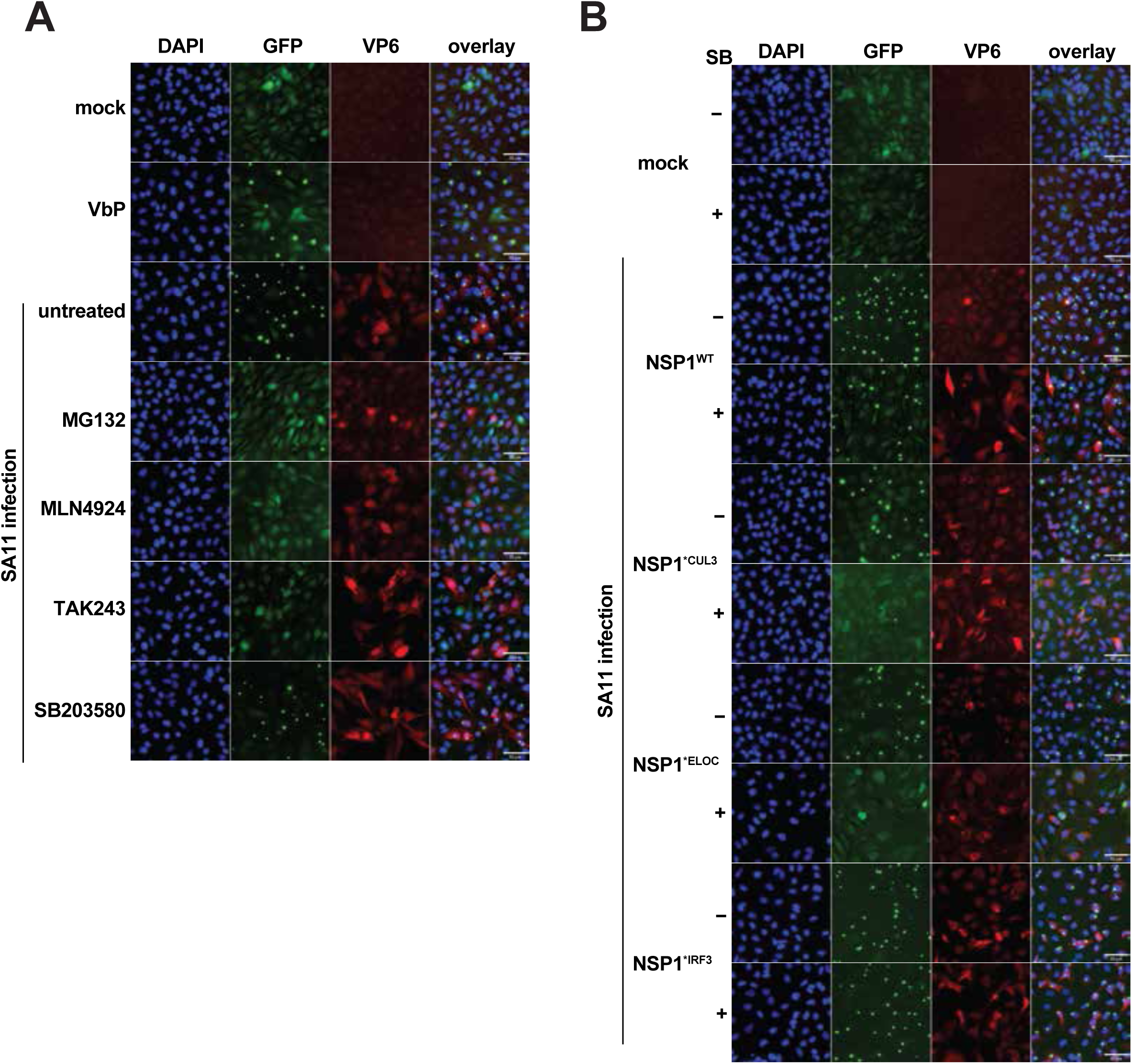
(A) Representative fluorescence microscopy images of A549-ASC-GFP-NLRP1 cells infected with SA11 (MOI=10, 8hpi), with or without treatment of SB202190 (20μM), MG132 (25μM), MLN4924 (10μM) or TAK-243 (5μM) post-inoculation. VbP (30μM) treatment served as a control for ASC activation. Images show representative fields. Scale bars, 50μm. (B) Representative fluorescence microscopy images of A549-ASC-GFP-NLRP1 cells infected with SA11-4F and its corresponding mutants SA11-4F-NSP1*^CUL3^, SA11-4F-NSP1*^ELOC^, or SA11-4F-NSP1*^IRF3^ (MOI=10, 8hpi), with or without treatment of SB202190 (20μM) 1h pre-inoculation. Images show representative fields. Scale bars, 50μm.

